# Early persistence of recipient stem-cells and T-cell dysregulation are associated with relapse after transplant in AML/MDS

**DOI:** 10.1101/2025.11.13.688350

**Authors:** Nurefsan E. Sariipek, Ksenia R. Safina, Wesley S. Lu, Shuqiang Li, J. Thomas Janes, McKayla Van Orden, Kenneth J. Livak, Haynes Heaton, Vincent T. Ho, John Koreth, Marlise R. Luskin, Gabriel K. Griffin, Jacqueline S. Garcia, R. Coleman Lindsley, Corey Cutler, Robert J. Soiffer, Jerome Ritz, Catherine J. Wu, Joseph H. Antin, Andrew A. Lane, Mahasweta Gooptu, Peter van Galen

## Abstract

Hematopoietic stem cell transplantation (HSCT) offers the best curative option for acute myeloid leukemia (AML) and myelodysplastic syndrome (MDS), yet relapse remains common. Current relapse detection methods are often too late for effective intervention. To identify earlier predictors and therapeutic targets, we performed longitudinal single-cell RNA and T cell receptor (TCR) sequencing of bone marrow from 33 AML/MDS patients during post-transplant immune reconstitution, comparing those who relapsed to those who remained in remission. Persistence of recipient hematopoietic stem and progenitor cells (HSPCs) in the marrow was associated with relapse months later. These residual recipient HSPCs harbored copy number variations (CNVs), supporting their leukemic origin, and overexpressed *PRAME* and *CALCRL* compared to coexisting donor HSPCs. Further, in a subset of *TP53*-mutant disease, low TCR diversity with skewing toward dominant clonotypes foreshadowed relapse. These findings lay the groundwork for improved relapse prediction and nominate therapeutic targets for early post-transplant intervention.

## INTRODUCTION

AML is one of the most challenging hematologic malignancies to treat, with a 5-year survival rate of 29%^1^. The standard therapeutic approach for intermediate and high-risk disease uses induction chemotherapy followed by consolidation in the form of allogeneic HSCT to eliminate remaining leukemia cells by means of immunologic killing, called the graft-versus-leukemia (GVL) effect. However, a common outcome of HSCT is disease relapse, which can occur in up to 80% of patients in high-risk genomic groups, such as those with *TP53* mutations^2–4^. Physicians can monitor blood and bone marrow for residual leukemia cells and changes in chimerism (the balance between donor and recipient cells). Reappearance of recipient mature hematopoietic cells in the peripheral blood, measured by standard peripheral blood leucocyte chimerism, often signals impeding relapse, but by then, treatment options are limited.

After HSCT, reconstitution of the blood system occurs through a progressive process. In the first three months, donor HSPCs undergo accelerated differentiation to reconstitute erythroid/megakaryocyte, myeloid, and eventually lymphoid lineages^5,6^. From three to six months, circulating immune cell composition is relatively stable, followed by decline in innate populations and an increase in adaptive populations before steady-state hematopoiesis is established (**Figure 1A**)^6^. The first three months post-transplant represent a critical window for T cells and NK cells to establish their repertoire^7^. These cells are particularly important for the GVL effect, which needs to be balanced with graft-versus-host disease (GVHD) to eliminate persistent AML cells while sparing healthy tissues^8,9^. The balance between effector and regulatory components of the adaptive immune system plays a critical role in determining post-transplant outcomes, although strategies to modulate this balance remain poorly defined.

**Figure 1.**
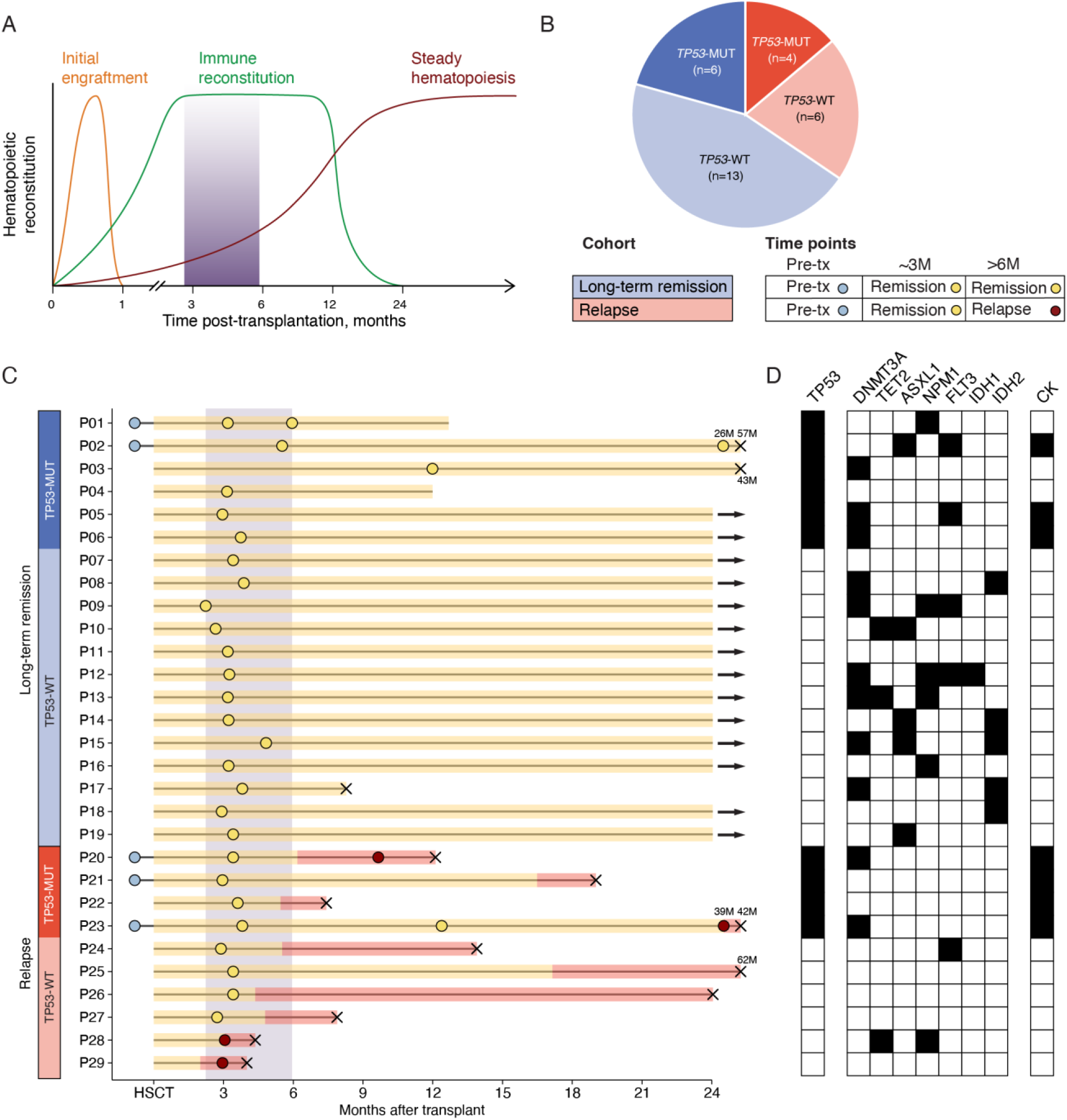
Cohort design for immune reconstitution early after transplant in AML/MDS patients. **A.** Schematic of hematopoietic reconstitution phases after transplant: initial engraftment (orange line), immune reconstitution (green line), and steady hematopoiesis (brown line). Much of our analysis focuses on bone marrow samples collected ∼3 months after transplant (shaded area). **B.** Pie chart shows sample characteristics: long-term remission cohort samples are in blue and relapse cohort in red, with darker shades indicating *TP53*-mutant and lighter shades indicating *TP53* wild-type. The table summarizes sample time points and disease stages: pre-transplant (light blue), remission (yellow), and relapse (red). **C.** Swimmer plot shows the clinical course of patients in our cohorts. Each horizontal bar represents one patient’s timeline. Circles mark time points of bone marrow sample collection. Yellow circles represent samples collected during remission; red circles during relapse, blue prior to HSCT. Bars shaded yellow indicate clinically remission period, red indicate relapse intervals. Black × denotes death. Arrows indicate patients who remained in remission beyond 42 months. The purple shaded region (∼3 months) marks the remission samples on which most analyses are focused. See **Supp. Figure 1A** for an extended version. **D.** Mutation plot for selected genes at diagnosis. Black squares indicate mutations corresponding to lines (patients) in panel C. CK: complex cytogenetics. See **Suppl. Table 1** for more details.

T cells have long been recognized as central mediators of the GVL effect, with CD8⁺ T cells and their receptors receiving particular attention. Over the years, advances in high-throughput sequencing have enabled characterization of the TCR repertoire, enabling accurate quantification of TCR diversity and their potential to recognize diverse antigens. After HSCT, TCR diversity may reflect the immune system’s capacity to recognize neoantigens and mount effective anti-leukemia responses^10–12^. Indeed, higher TCR diversity has been associated with improved clinical outcomes of patients with hematological malignancies receiving different transplant types^13–16^. These studies measured TCR diversity in the peripheral blood and did not account for disease genetics, and effect sizes were modest. Thus, the importance of TCR repertoire reconstitution remains unclear in the bone marrow, the primary site of disease.

Current tools remain limited in their ability to identify patients at risk for relapse. In the clinic, post-HSCT chimerism is routinely assessed using DNA-based methods that quantify recipient and donor cell proportions. While most patients maintain high donor chimerism under standard monitoring, a decline—which typically precedes morphologic or molecular relapse by only days to weeks—usually occurs too late for pre-emptive intervention. Earlier changes in the immune landscape may signal relapse, as suggested by expression of inhibitory receptors on CD4 T cells before relapse^6^ and downregulation of antigen presentation on AML blasts at relapse^17,18^. More granular insight into cellular and molecular dynamics in complex tissues can now be obtained using single-cell technologies^19^, exemplified by the recent identification of a CD8+ T cell subset associated with effective anti-leukemia responses^20^. Developments in single-cell genomics now allow for the acquisition of multiple layers of information in addition to gene expression, including copy number variations (CNVs), genetic variation that distinguishes donor and recipient cells, and TCR sequences^21–24^. By integrating these modalities, it is now within reach to characterize rare cell types that drive leukemia progression and their interaction with the immune system.

To this end, we studied immune reconstitution dynamics in 49 longitudinal bone marrow samples from 33 patients undergoing HSCT. We focused our analysis on early time points, ∼3 months (range 2.2-5.9 months) after transplant, while patients were still in morphologic remission with intact peripheral blood chimerism, to identify features that might be used to predict and/or prevent relapse. A strong association with relapse was the persistence of recipient HSPCs, which harbored leukemic CNVs. By comparing these recipient HSPCs with co-existing donor HSPCs in the same environment, we identified potential immunotherapeutic targets to specifically eliminate the former. Furthermore, within the high-risk subgroup of patients with *TP53*-mutated disease, low TCR diversity foreshadowed relapse. The identification of relapse-associated features lays the groundwork for improved risk stratification and more effective post-transplant interventions.

## RESULTS

### Cohort design to study immune reconstitution early after transplant in AML/MDS

To investigate the immune landscape following stem-cell transplantation, we analyzed 29 AML/MDS patients who achieved clinical remission before undergoing T cell-replete HSCT. All patients received tacrolimus-based prophylaxis except one who received post-transplant cyclophosphamide-based prophylaxis. Patients were stratified into two cohorts: those who remained relapse-free during follow-up (n=19, median follow-up 36 months, range 8–83 months) and those who experienced relapse (n=10, median time to relapse 5.5 months, range 2–39 months). Four additional patients were only included for relapse analysis below and had measurable blasts at transplant, bringing the total to 33 patients (**Figure 1B-C, Suppl. Figure 1a**).

We collected bone marrow samples from multiple time points, including pre-transplant, during morphologic remission around 3 months, long-term remission, and—when applicable—at relapse. We focused our analysis on remission samples collected ∼3 months after transplant (n=27, median 3.3 months, range 2.2–5.9 months), the earliest time when GVL activity is substantially established and early therapeutic intervention may prevent later relapse. Unless otherwise noted, all analyses were restricted to remission samples collected at this ∼3-month timepoint.

The patients in the study harbored various genetic alterations. Given that the clinical outcome of relapse post-transplant is particularly germane in *TP53-*mutated AML with relapse rates in the 80% range^2,3^, we included *TP53*-mutated AML in both long-term remission (n=6 of which three were *TP53* multi-hit) and relapse cohorts (n=4, all *TP53* multi-hit) (**Figure 1B-C**). In addition, patients harbored recurrent mutations including *DNMT3A* (n=11), *NPM1* (n=6), *IDH2* (n=5), and *ASXL1* (n=5) (**Figure 1D, Suppl. Table 1**).

### Skewed T cell reconstitution precedes relapse in *TP53*-mutated AML/MDS

We performed paired single-cell RNA sequencing and TCR sequencing on 49 samples from 33 patients in total, yielding 496,691 high-quality single cells (**Figure 2A**). For cell type annotation, we used the recently published AML-specific bone marrow reference atlas, which enabled efficient and accurate classification of hematopoietic and immune cell states^25^. Inspection of resulting cell type-specific gene expression confirmed that the 35 cell types expressed canonical marker genes (**Suppl. Figure 1B**).

**Figure 2.**
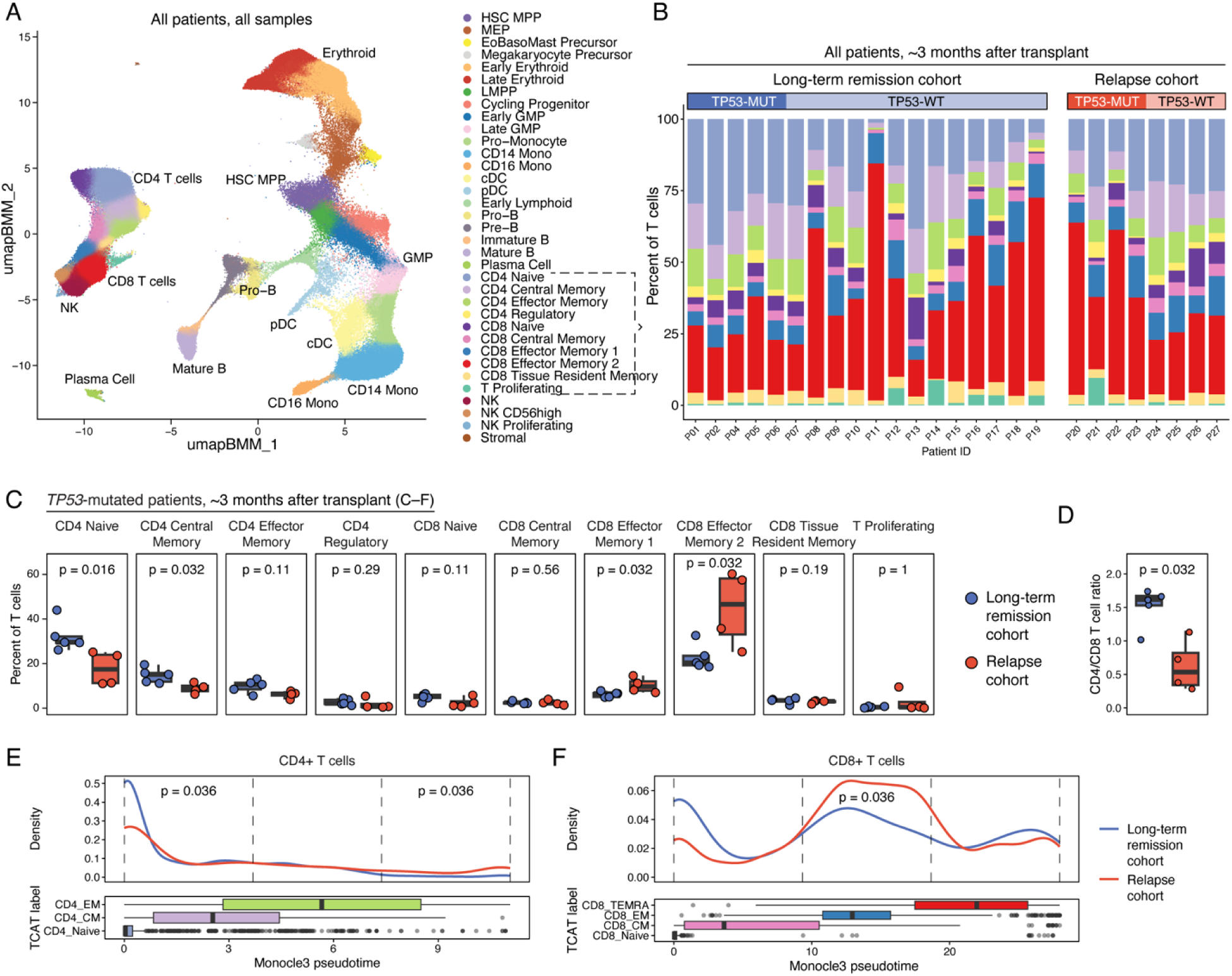
Skewed T cell reconstitution precedes relapse in patients with *TP53*-mutated AML/MDS. **A.** UMAP projection of 496,691 single-cell transcriptomes from 49 bone marrow samples across 33 patients, colored by cell type. Corresponding annotations are shown on the right. **B.** Stacked bar plot depicts T cell subset proportions in patients from the long-term remission (left) and relapse (right) cohorts. Each bar represents a sample collected ∼3 months after transplant remission. No significant differences in T cell composition were observed between the full cohorts. **C.** Boxplots comparing the proportion of T cell subsets between long-term remission and relapse cohorts among patients with *TP53*-mutated AML/MDS ∼3 months after transplant. CD4 Naive and CD4 Central Memory cells are significantly more abundant in the remission group, while CD8 Effector Memory 1 and CD8 Effector Memory 2 cells are significantly enriched in the relapse group. P-value was calculated using Wilcoxon test. **D.** Comparison of CD4/CD8 ratio in patients with *TP53*-mutated AML/MDS ∼3 months after transplant, showing significantly reduced levels in the relapse cohort P-value was calculated using Wilcoxon test. **E-F.** Top: Density plots show the distribution of Monocle3-inferred pseudotime values for (**E**) CD4⁺ and (**F**) CD8⁺ T cells. The relapse cohort (red line) exhibits a rightward shift in pseudotime, suggesting more differentiated T cell states compared to the long-term remission cohort (blue line). Bottom boxplots show the pseudotime distribution of annotated T cell subsets (TCAT annotations). Vertical dashed lines indicate three pseudotime bins used to compare the proportion of cells between cohorts using a Wilcoxon test (**Suppl. Figure 2F**). BoneMarrowMap labels: HSC: hematopoietic stem cells, MPP: multipotent progenitors, MEP: megakaryocyte-erythrocyte progenitors, EoBasoMast precursor: eosinophil/basophil/mast cell precursors, LMPP: lymphoid-primed multipotent progenitors, GMP: granulocyte-monocyte progenitors, Mono: monocytes, cDC: conventional dendritic cells, pDC: plasmacytoid dendritic cells, CD4 CM: CD4^+^ central memory, CD4 EM: CD4^+^ effector memory, CD8 TEMRA: CD8^+^ effector memory-expressing CD45RA, CD8 EM: CD8^+^ effector memory, CD8 CM: CD8^+^ central memory.

We first compared the distribution of T cell subtypes ∼3 months after transplant since this is a critical time for post-transplant immune reconstitution. However, we did not observe significant differences between the cohorts, i.e., patients who remain in long-term remission and patients who would later develop relapse (**Figure 2B, Suppl. Figure 2A-B**). We next focused on patients with *TP53*-mutated AML, which is the highest-risk subset, ∼3 months after transplant. Here, we found significantly lower levels of naïve CD4+ T cells in patients who eventually relapsed compared to those who remained in long-term remission (**Figure 2C**). In addition, the relapse cohort demonstrated significant increases in CD8+ effector memory T cell subsets (EM1 and EM2). We also examined the CD4/CD8 ratio, a broad marker of immune health in infections and cancer^26,27^. In patients with *TP53*-mutated AML who remained in long-term remission, the median CD4/CD8 ratio was 1.62 (**Figure 2D**), resembling levels seen in healthy individuals. In contrast, patients who later relapsed had significantly lower CD4/CD8 ratios (median=0.54, p=0.032). The patients with *TP53*-mutated AML/MDS were in morphologic remission at this time and did not develop relapse until a median of 8.1 months after sampling (range 1.8–35.6 months). These findings suggest that early immune dysregulation, reflected in T cell composition, may precede relapse in patients with *TP53*-mutated AML/MDS.

To further dissect T cell differences between cohorts, we used T-CellAnnoTator (TCAT), a tool designed for refined T cell annotation and functional profiling from single-cell RNA-seq data^28^. While we confirmed the lower CD4/CD8 ratio in patients with *TP53-*mutated AML in the relapse cohort compared to the long-term remission cohort, we did not observe statistically significant differences in gene expression program activity within T cell subsets (**Suppl. Figure 2C**). Next, TCAT’s more granular annotations allowed us to select specific T cell subsets for pseudotime analysis–a computational approach that orders cells along a trajectory, providing a proxy for their progression through gene expression states. We selected naïve (primitive), memory (intermediate), and effector (terminal) CD4+ and CD8+ T cells and applied the pseudotime algorithm Monocle3 to reconstruct T cell trajectories (**Suppl. Figure 2D-E**). We then binned CD4+ and CD8+ in three bins of low, medium, and high pseudotime as a proxy for progressive differentiation states. Comparing the distribution of T cells across these states, we found a shift towards higher pseudotime bins in patients with *TP53*-mutated AML who eventually relapsed compared to those who remained in long-term remission (**Figure 2E, Suppl. Figure 2F**). Mirroring the T cell proportion analysis that showed decreased CD4+ naïve T cells and increased CD8+ effector cells, these data suggest a link between accelerated T cell differentiation and a higher risk of relapse in patients with *TP53-*mutated AML/MDS.

### Low TCR diversity precedes relapse in *TP53*-mutated AML/MDS

To further explore the implications of early T cell differences, we examined TCR clonality in samples collected ∼3 months after transplant. We captured TCR sequences in 85.6% of all 199,957 T cells in our dataset, representing 115,205 unique clonotypes (defined by a shared TCR sequence, **Figure 3A**, **Suppl. Figure 3A**). CD8+ T cells harbored larger clones than CD4+ T cells, and clone size increased in more differentiated subsets (**Figure 3B**). When comparing clone sizes between long-term remission and relapse cohorts ∼3 months after transplant, we found no significant difference, though when limited to patients with *TP53*-mutated AML/MDS, we observed a trend towards fewer, more expanded clonotypes in relapse-fated patients (**Figure 3B, Suppl. Figure 3B-C**).

**Figure 3.**
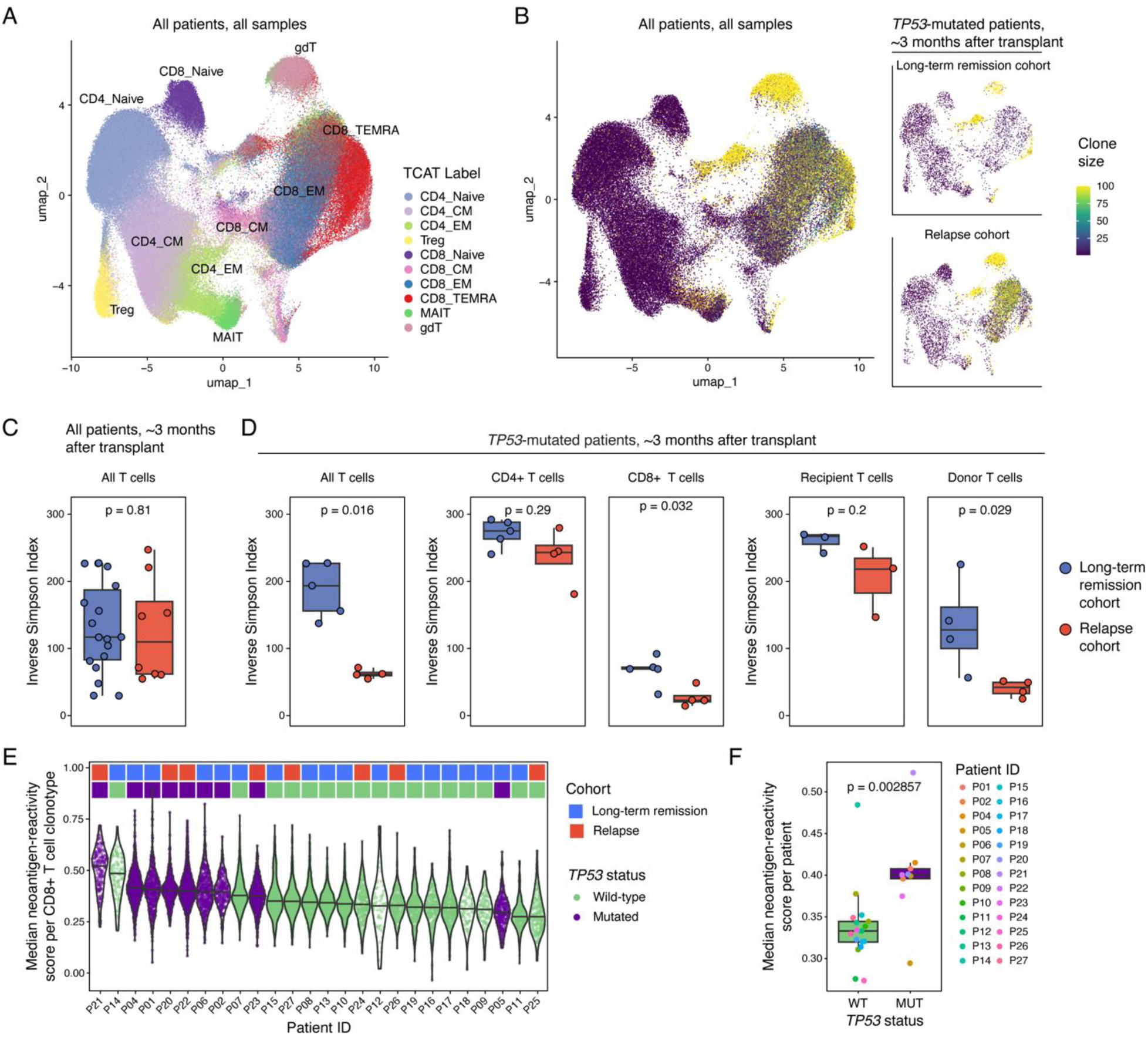
Low TCR diversity precedes relapse in *TP53*-mutant AML/MDS, which shows higher antigen-recognition signatures than *TP53* wild-type disease. **A.** UMAP projection of all T cells from all samples, color-coded by T cell subset labels from TCAT. **B.** UMAP shows T cells colored by clone size. The inset panels show clone size in samples from patients with *TP53*-mutated AML ∼3 months after transplant, separated by long-term remission and relapse cohorts. **C.** Boxplot shows TCR diversity indices in all samples that were collected ∼3 months after transplant in remission, separated by cohort. All samples were subsetted to 300 T cells. P-value was calculated using Wilcoxon test. **D.** Boxplots show TCR diversity indices in the same samples, subsetted for patients with *TP53*-mutated AML. Data is shown for all T cells (left), stratified by T cell subset (CD4+ or CD8+), and stratified by recipient vs. donor origin. All samples were subsetted to 300 T cells or excluded if fewer T cells were detected. P-values were calculated using Wilcoxon tests. **E.** Violin plot shows CD8+ neoantigen-reactivity scores in CD8+ T cells ∼3 months after transplant. Each symbol represents a T cell clonotype (defined by shared TCR sequence), with values indicating the median NeoTCR8 score^29^ across all cells within that clonotype. **F.** Boxplot shows median CD8+ neoantigen-reactivity scores in CD8+ T cells ∼3 months after transplant per patient, separated by *TP53* status. Each symbol represents the median score of all T cell clonotypes within one patient. P-value was calculated using Wilcoxon test. TCAT labels: CD4_CM: CD4 Central Memory, CD4_EM: CD4 Effector Memory, Treg: T regulatory cell, CD8_CM: CD8 Central Memory, CD8_EM: CD8 Effector Memory, CD8_TEMRA: CD8 T Effector Memory-Expressing CD45RA, MAIT: Mucosal-associated invariant T cells (MAIT), and gdT: γδ T cells.

TCR diversity, as described by the inverse Simpson index, provides a metric to estimate the evenness of clonotype distributions within a T cell repertoire. We calculated the inverse Simpson index, in which a higher value indicates greater TCR diversity, for all patients ∼3 months after transplant. When comparing the full long-term remission and relapse cohorts, there was no significant difference in TCR diversity (**Figure 3C, Suppl. Figure 3D**). However, when restricting our analysis to *TP53*-mutated samples, we found that patients who later relapsed had lower T cell diversity compared to patients who stayed in long-term remission (**Figure 3D**). This difference was primarily driven by donor CD8+ T cells (**Figure 3D**), which we distinguished from recipient T cells using the Souporcell package as detailed below^22^. Together with differentiation differences noted above, reduced TCR diversity indicates that disordered T cell reconstitution distinguishes relapse from remission in *TP53*-mutated AML/MDS and may serve as an indicator of relapse risk.

### CD8+ T cell clones show higher neoantigen reactivity scores in *TP53*-mutant versus *TP53*-wildtype disease

Motivated by the T cell dynamics associated with outcome in *TP53*-mutated patients, and considering the role of antigen-specific T cell responses in anti-tumor immunity, we wanted to investigate gene expression signatures associated with antigen recognition. To this end, we applied neoantigen-reactivity signatures from the Rosenberg lab that incorporate transcriptional features of T cell recognition of tumor-specific neoantigens^29^. We scored expression of this signature in each T cell ∼3 months after transplant and then ranked T cell clonotypes by their median neoantigen-reactivity score. These scores were not associated with long-term remission vs. relapse cohorts (**Figure 3E**, top bar). However, we found that patients with *TP53*-mutated AML had higher scores compared to patients with *TP53* wild-type AML (fold change 1.20, p = 0.003, **Figure 3E-F**). To confirm this finding, we also calculated the antigen-specific activation (ASA) score using TCAT, which sums four activation associated gene expression programs as a proxy for TCR activation^28^. This analysis confirmed that CD8+ clonotypes from patients with *TP53*-mutated AML exhibited elevated antigen recognition scores (fold change 1.31, p = 0.045, **Suppl. Figure 3E-F**). CD4+ T cells did not show a significant difference in neoantigen-reactivity scores (**Suppl. Figure 3G-H**). Despite the higher antigen recognition scores in CD8+ T cells from patients with *TP53*-mutated AML compared to *TP53* wild-type AML, their high rates of relapse suggest ineffective GVL and leukemic cell clearance.

### Recipient cell persistence in the stem and progenitor cell compartment is associated with relapse

Next, we shifted our focus beyond T cells to analyze the hematopoietic system more broadly. To quantify immune reconstitution dynamics with more precision, we wanted to distinguish recipient- and donor-derived cells from each other. To this end, we applied Souporcell, an algorithm that uses natural genetic variation to infer cell genotypes directly from single-cell RNA-seq data without prior genotype information (**Figure 4A**)^22^. Specifically, each cell is assigned as recipient or donor based on its expression of reference versus alternative alleles, which can then be analyzed in combination with metadata such as time point and cell type (**Figure 4B**). We developed a set of rules to select patients with reliable recipient and donor cell calls (see **Methods**), resulting in successful resolution of 21 of the 33 patients, yielding recipient/donor assignments for 342,780 out of 496,691 cells.

**Figure 4.**
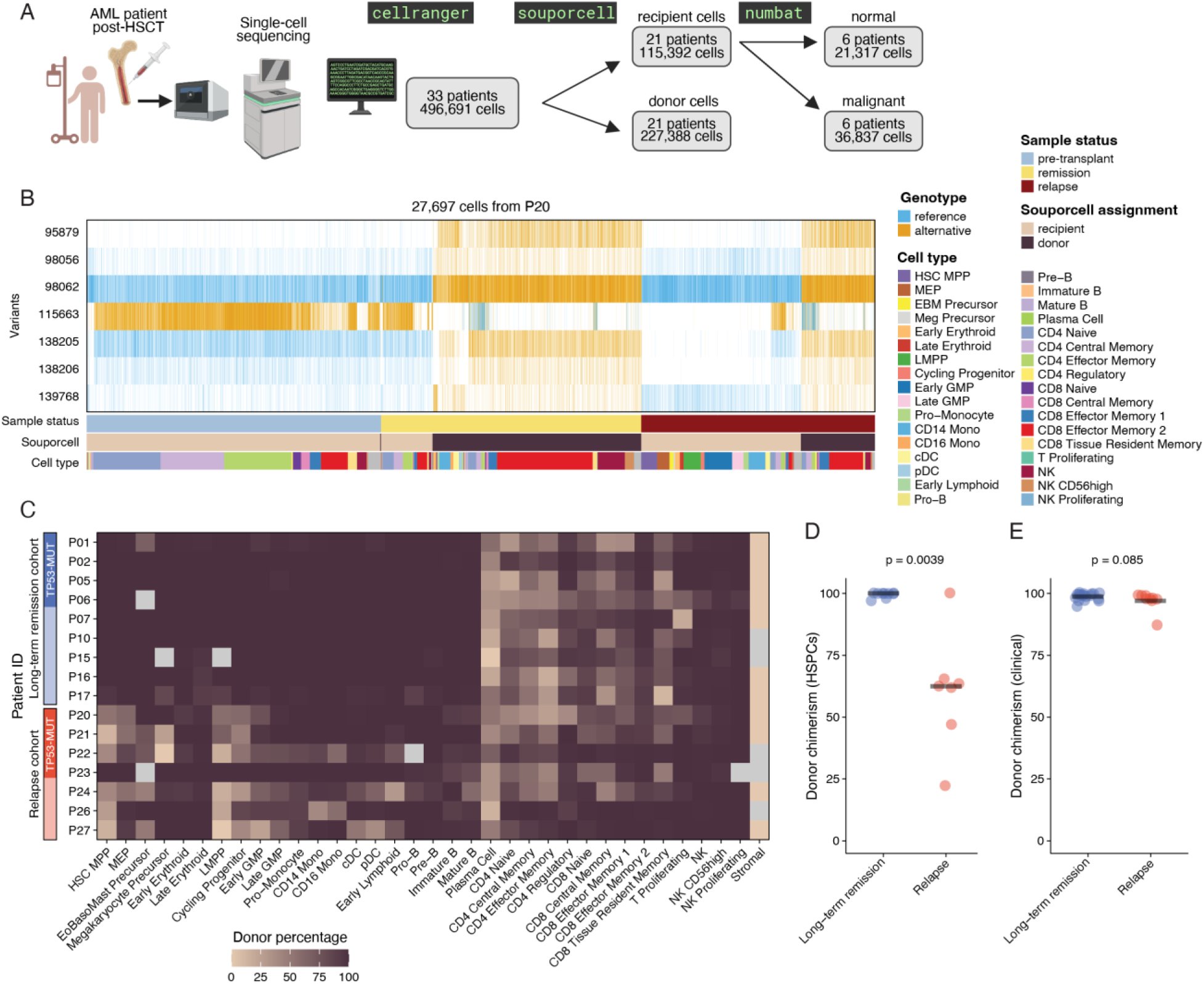
Recipient HSPC persistence is an early predictor of relapse. **A.** Overview of experimental and computational pipeline. Single-cell sequencing data from AML/MDS patient bone marrow samples was processed by Cell Ranger, followed by genotype-based cell origin assignment using Souporcell (to categorize cells into recipient- and donor-derived) and CNV-based malignancy assessment using Numbat (to categorize cells into malignant and normal). **B.** Example Souporcell variant heatmap from patient P20 shows variant allele patterns used for donor vs. recipient assignment. Each column represents a cell, each row represents a selected variant. Cell annotations at the bottom include sample status (pre-transplant, remission, relapse), Souporcell assignment, and cell type. **C.** Heatmap shows the proportion of donor-derived cells across different cell types and patients from remission samples collected ∼3 months after transplant. Each row represents a patient, grouped by long-term remission or relapse cohort, and each column corresponds to a cell type. Color ranges from beige (low donor contribution; higher recipient-derived cells) to purple (high donor contribution). Grey indicates 0 cells were detected. **D.** Donor chimerism in HSPCs (HSCs, MPPs, MEPs, LMPPs, early GMPs, and cycling progenitors) at the remission time point ∼3 months after transplant. Patients in the relapse cohort (red, n=7) exhibit significantly lower donor chimerism compared to those in the long-term remission cohort (blue, n=9), months before relapse occurs. *P*-value was calculated using the Wilcoxon test. **E.** Clinically assessed donor chimerism at ∼3 months after transplant shows no significant difference between the long-term remission (blue, n=18) and relapse (red, n=9) cohorts. *P*-value was calculated using the Wilcoxon test.

Leveraging this genotype-based chimerism analysis, we examined the proportions of recipient- and donor-derived cell types in remission samples ∼3 months after transplant. As expected, the replacement of recipient cells by donor cells varied across cell populations, with the persistence of recipient cells being highest for stromal cells and plasma cells (100% and 59%, respectively, **Figure 4C, Suppl. Figure 4A**). The persistence of recipient stem and progenitor cells appeared to be higher in patients who would later develop relapse compared to the long-term remission cohort. The exception was Patient 23, who relapsed 3.3 years after transplant, significantly later than others in the relapse cohort. This patient’s reconstitution dynamics more closely resembled those of the long-term remission cohort (**Figure 4C**). These observations prompted us to formally test if there was a difference in stem and progenitor chimerism between the cohorts.

To obtain larger cell numbers for chimerism estimates, we merged six progenitor populations (HSCs, MPPs, MEPs, LMPPs, early GMPs, and cycling progenitors) based on transcriptional similarity and cluster proximity (**Figure 2A**). We next quantified chimerism in this merged HSPC compartment ∼3 months after transplant. HSPC donor chimerism was significantly lower in patients who eventually relapsed compared to those who stayed in remission (median 63% and 100%, p=0.0039) (**Figure 4D, Suppl. Figure 4B**). In contrast, clinically used donor chimerism measurements in whole blood, that evaluate mature leucocytes rather than HSPCs, did not predict relapse (median 99% and 98%, p=0.085, **Figure 4E**). All 10 patients with <5% recipient HSPCs ∼3 months after transplant remained in remission for more than three years (remission cohort and patient 23), whereas all six patients with >5% developed relapse within 18 months (median 5.5, range 4.4-16.5 months). These six patients did not develop relapse until a median of 2.3 months after sampling (range 1.0–13.5 months). This highlights the potential of HSPC chimerism for predicting relapse risk months before relapse occurs. Further, it suggests that persistence of recipient HSPCs in the bone marrow niche may be a key event leading to eventual relapse post-transplant.

### Recipient HSPCs persisting in remission are leukemic in origin

Next, we wanted to investigate what proportion of persistent recipient cells are part of the malignant clone (as opposed to residual normal hematopoiesis). For this, we used Numbat, a haplotype-aware CNV caller for single-cell sequencing data (**Figure 4A**)^23^. Numbat combines information from gene expression, allelic ratios, and population-derived haplotypes to detect allele-specific CNVs in single cells and to map their lineage relationships. Out of the 21 patients with Souporcell recipient cell annotations, Numbat identified CNVs in six patients. The CNV calls were consistent with clinical cytogenetics, such as deletions on chr5 and chr7 in Patient 33 (**Figure 5A, Suppl. Figure 5**). Based on CNV detection, Numbat assigned 21,317 normal and 36,837 malignant cells for the six patients across time points.

**Figure 5.**
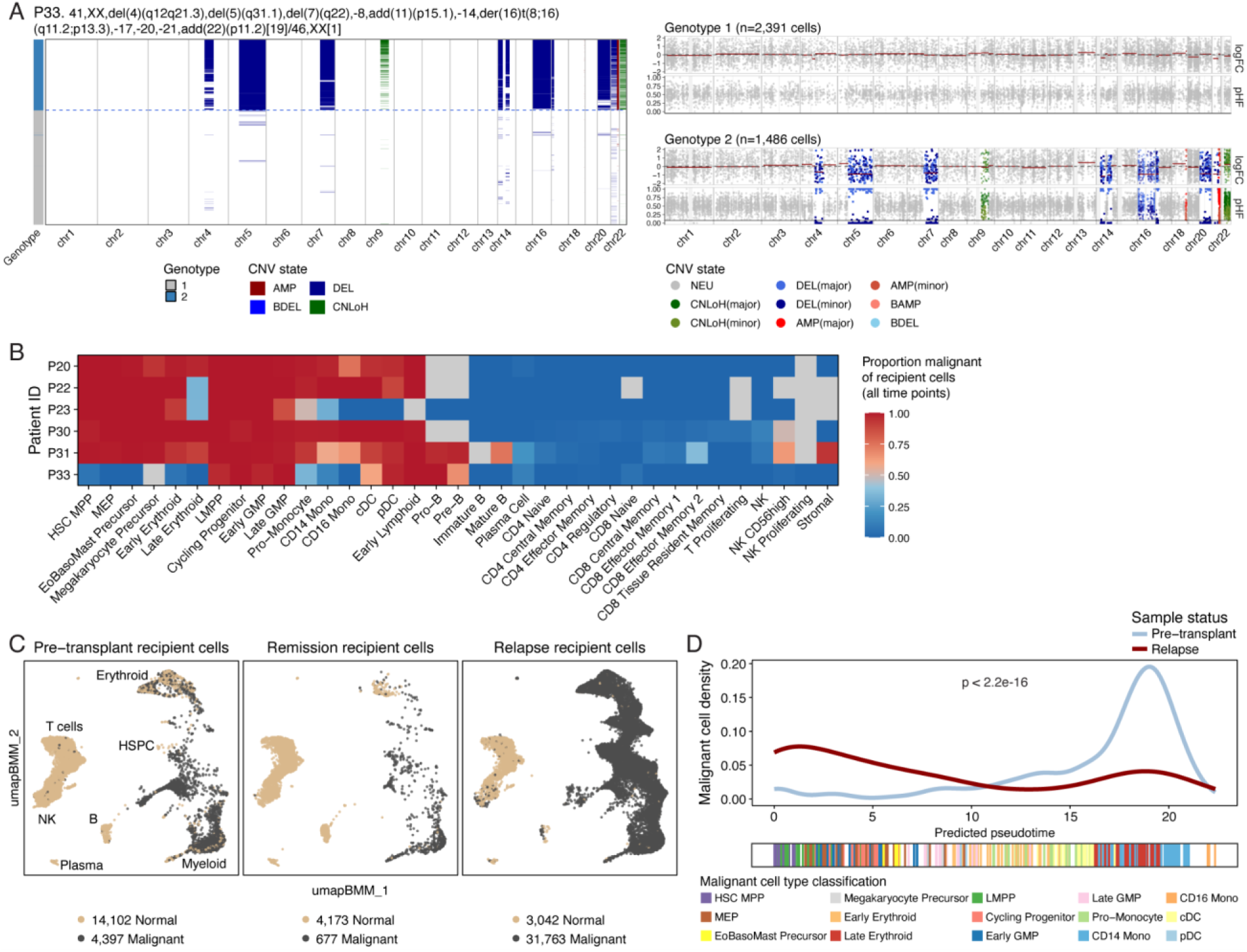
Malignant cells become more primitive over the course of disease progression. **A.** Left: heatmap shows genome-wide CNVs (colors) in single cells (rows) for Patient 33. Cytogenetics determined by karyotyping are indicated above the plot. Each row represents a cell, and columns correspond to genomic positions across chromosomes 1–22. CNV states are color-coded as indicated below the heatmaps: amplifications (AMP, dark red), deletions (DEL, dark blue), biallelic deletions (BDEL, light blue), and copy-neutral loss of heterozygosity (CNLoH, green). On the left, cells are grouped by genotype: normal cells (grey) and malignant cells (blue), separated by the horizontal dashed line. Right: genome plots show CNV profiles for pseudobulked clones, with pseudobulked normal cells on top (genotype 1) and pseudobulked malignant cells on the bottom (genotype 2). Each symbol represents a genomic bin, with read coverage (logFC, −2 to +2) shown on top and posterior haplotype frequencies (pHF, 0 to 1) on the bottom. **B.** Heatmap shows the proportion of malignant cells across six patients for whom CNV information was available, as inferred from Numbat analysis. Rows represent individual patients, and columns represent annotated cell types. Color indicates the proportion of malignant cells, ranging from blue (normal) to red (malignant). Grey indicates 0 cells were detected. Cells from all time points are included and P33 mainly represents pre-transplant cells. **C.** UMAP projection using the same embedding as in **Figure 2**, restricted to recipient cells from the six patients with Numbat-derived CNV calls. Cells are colored by Numbat compartment and plots are separated by sampling time point. **D.** Density plot shows the distribution of predicted pseudotime values for malignant cells from pre-transplant and relapse samples. Pseudotime and cell type annotations were determined by mapping cells onto the BoneMarrowMap reference. The bottom bar shows malignant cells colored by cell type classification to illustrate that pseudotime values are a proxy for differentiation. Statistical significance was assessed using the Kolmogorov–Smirnov test.

The proportion of recipient cells harboring CNVs differed between cell types (**Figure 5B**). Most cells in the HSPC, erythroid, and myeloid lineages were classified as malignant, whereas most B, T, and NK cells were not (**Figure 5B**). The presence of CNVs in a subset of residual recipient cells supports their malignant identity. Notably, for all patients where Numbat detected CNVs and we analyzed a remission sample ∼3 months after transplant, every recipient HSPC in remission harbored CNVs (75 cells across 3 patients, **Suppl. Fig. 5B**), establishing these persistent HSPCs as leukemic in origin.

### Malignant cells shift towards primitive states at relapse

Out of the six patients in which Numbat identified CNVs, four had sufficient malignant cells pre-transplant and at relapse to evaluate malignant cell evolution over time (n=2,911 and n=31,763 cells, **Suppl. Figure 6A**). Between pre-transplant and relapse malignant cells, 1,414 genes were differentially expressed (**Suppl. Figure 6B-D**). We first examined MHC class II, which has been proposed as a key mechanism of immune evasion following allogeneic transplantation^17,18^. Surprisingly, we did not observe a significant decrease in expression of MHC-II members in relapse compared to pre-transplant cells in this cohort of mainly fully-matched donor transplants (**Suppl. Table 2**). To learn more about pathways that are altered during disease progression, we performed gene set enrichment analysis (GSEA). This showed that malignant cells at relapse, compared to pre-transplant, upregulate HSC-associated gene signatures (**Suppl. Figure 6E**). This prompted us to investigate the developmental trajectory of AML/MDS cells during disease progression. Using pseudotime analysis based on the bone marrow reference atlas^25^, we placed tumor cells along a continuum of hematopoietic differentiation. Relapse tumor cells appeared less differentiated compared to pre-transplant tumor cells (**Figure 5D**), consistent with GSEA showing enrichment of stemness-associated signatures. This effect was observed in each of the four patients (**Suppl. Figure 6F**). These findings indicate a shift toward less differentiated cell states from pre-transplant to relapse, raising questions about the properties of residual recipient HSPCs during remission.

### Persistent recipient HSPCs show distinct antigenic profiles and reduced proliferation

Our strategy to annotate cell origins provides a first-of-its-kind opportunity to compare recipient HSPCs to donor HSPCs at remission (**Figure 6A**). For this analysis, we excluded the long-term remission cohort which had <10 persistent recipient HSPCs and P30-P33 who did not have remission samples at the 3-month time point. Thus, the following analysis of recipient vs. donor HSPCs during remission is based on samples from six patients who developed relapse within 18 months (**Suppl. Figure 7A**). Persistent recipient HSPCs made up 0.27–1.31% of these samples and, as noted above, CNV analysis indicated they are leukemic in origin (**Suppl. Figure 5B).**

**Figure 6.**
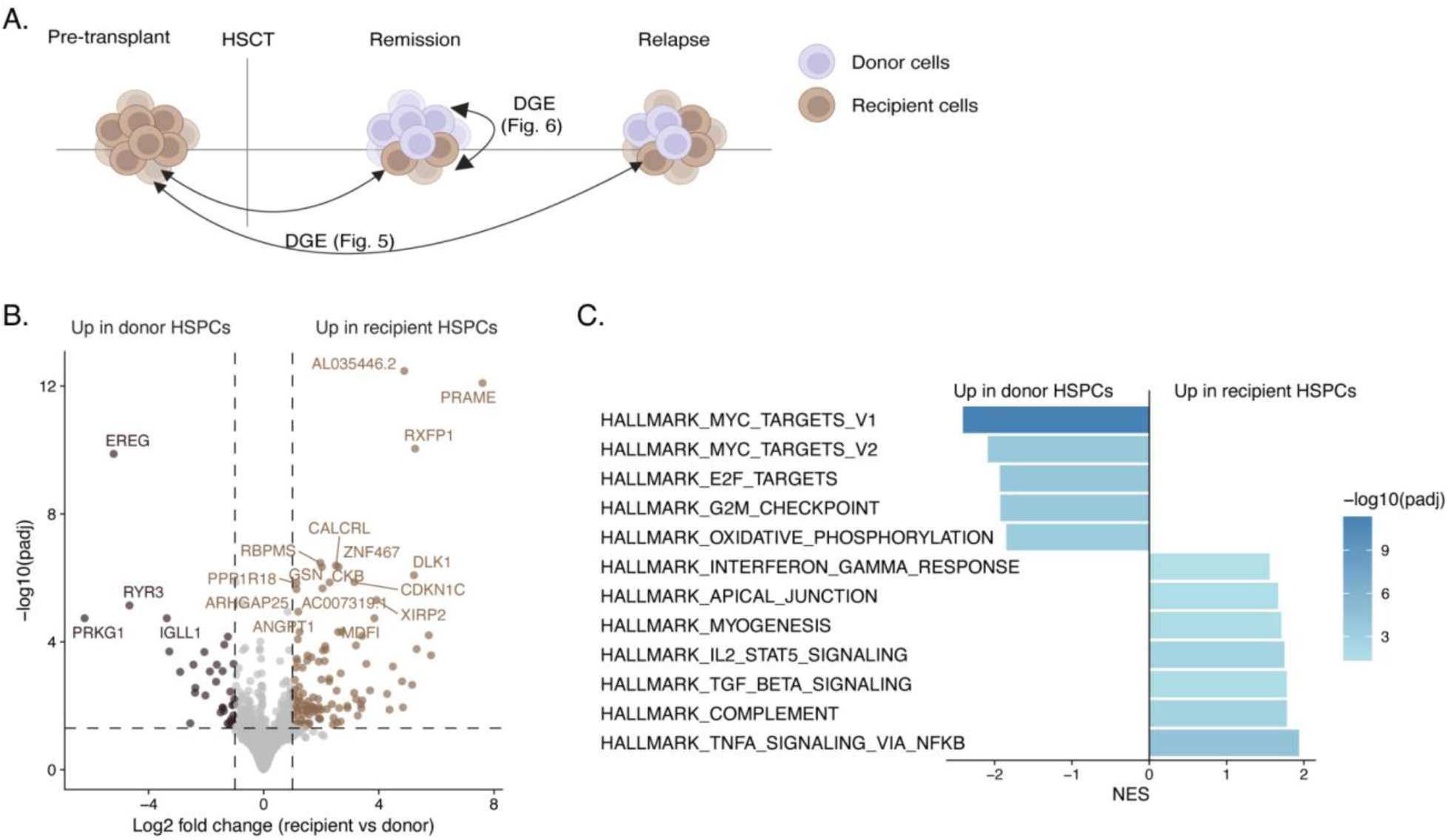
Persistent recipient HSPCs in remission have a distinct antigenic profile and reduced proliferation. **A.** Diagram depicts differential gene expression (DGE) comparisons made between pre-transplant and relapse cells (in **Figure 5** and **Suppl. Figure 6**) and between recipient and donor HSPCs in remission samples ∼3 months after transplant (in Figure 6 and **Suppl. Figure 7**). **B.** Volcano plot shows differentially expressed genes (symbols) between recipient HSPCs (n=533 cells across six patients) and donor HSPCs (n=540 cells across the same six patients) during remission. **C.** Bar plot shows pathways that were enriched (right) or depleted (left) in recipient HSPCs compared to donor HSPCs by GSEA. NES: normalized enrichment score.

In recipient HSPCs (n=533 cells) compared their donor counterparts (n=540), 118 genes were upregulated and 36 were downregulated (**Figure 6B, Suppl. Figure 7B-C, Suppl. Table 3**). Upregulated genes might confer a therapeutic window, and included potentially targetable tumor-associated antigens (*PRAME, CALCRL, RXFP1*) and a leukemia stem cell quiescence-associated gene (*CDKN1C*)—discussed in more detail below. To further characterize the transcriptional state of recipient HSPCs, we performed GSEA using the Hallmark pathways. This analysis revealed downregulation of cell cycle-related pathways in persistent recipient HSPCs and upregulation of immune signaling such as IL-2/STAT5 signaling, TGF-β signaling, and TNF-α signaling via NF-κB (**Figure 6C**). Thus, our comparison of recipient vs. donor HSPCs from the same bone marrow in remission highlights features that might be specifically targeted to prevent relapse.

## DISCUSSION

Our assessment of longitudinal samples from patients undergoing HSCT revealed several features of early immune reconstitution that are associated with subsequent relapse. The persistence of recipient HSPCs in the bone marrow niche identified patients at risk of relapse with high specificity. Comparing malignant recipient HSPCs with co-existing donor HSPCs in the same marrow revealed high expression of potential immunotherapeutic targets in the former. Further, within the subgroup of patients with *TP53*-mutant AML/MDS, low CD4/CD8 ratios and low TCR diversity foreshadowed relapse, suggesting that impaired immune surveillance may be associated with poor outcomes in this genetic context. By providing a deeper understanding of the cellular components and immune dynamics after HSCT, our study reveals strategies for relapse prediction and potential targets for early intervention.

Current post-transplant surveillance tools, such as traditional leucocyte chimerism analysis, often fail to provide actionable lead time, with changes typically emerging only days to weeks before clinical relapse. Given the dismal prognosis associated with post-transplant relapse, earlier predictors could enable timely, risk-adapted interventions. Around three months after transplant, conventional assays measured donor chimerism at 99% for patients who remained in remission and 98% for relapse-fated patients, whereas our assay measured 100% and 63% donor HSPCs, respectively. This measurement preceded relapse detection by more than two months, providing a potential window for intervention. The resolution to measure chimerism in specific cell types was afforded by using genetic variation to classify recipient and donor cells while concurrently measuring single-cell transcriptomes. While clinical implementation of single-cell sequencing may be challenging, these results should motivate development of chimerism assays focused on bone marrow HSPCs rather than whole blood leucocytes, which may be achievable with low-input chimerism assays^30^.

In the context of *TP53* mutations, a high-risk subgroup of patients with up to 80% relapse after transplant, T cell analysis afforded additional discriminatory value for relapse prediction. Several T cell properties differed between patients with *TP53*-mutated AML/MDS who later developed relapse and those who remained relapse-free. Relapse-fated patients had lower naïve and higher effector T cell proportions, suggesting accelerated differentiation. They also had low CD4/CD8 ratios and low TCR diversity with a tendency towards clonotype expansion. The association between low TCR diversity and relapse is supported by prior studies that evaluated peripheral blood in various hematological malignancies and transplant types (chiefly T-cell depleted transplants)^13–15^. With the development of reliable TCR sequencing methods from limited DNA, this may serve as a useful prognostic indicator.

The differences in T cell proportions and TCR diversity were restricted to *TP53*-mutated AML/MDS. It is not unexpected that this disease subset has a unique immune environment. *TP53* mutations affect cancer–immune interactions, including resistance to immune-mediated killing and increased secretion of cytokines and chemokines that impair NK- and T-cell–mediated anti-tumor immunity^31–35^. Our gene expression analysis indicates higher antigen recognition by CD8+ T cell clonotypes in patients with *TP53*-mutated AML/MDS compared to *TP53* wild-type AML, but this antigen recognition may be followed by an ineffective GVL response, given the poor outcome of patients with *TP53*-mutated disease. Although the mechanisms by which *TP53*-mutated AML uniquely evades anti-tumor immunity are beyond the scope of this study, defining which of the >100,000 TCR clonotypes we recovered recognize AML blast antigens will be an important next step.

In line with studies of AML under chemotherapeutic pressure, disease evolution from pre-transplant to relapse involved a shift toward a more primitive state, consistent with selection during conditioning, transplantation, or immune surveillance^36–38^. Further insight into cancer cell adaptation comes from our analysis of recipient HSPCs, which are likely relapse precursors as they harbor malignancy-associated CNVs. Compared with donor HSPCs competing in the same remission bone marrows, recipient HSPCs downregulated proliferation-related signatures, possibly reflecting prior cytotoxic therapy, and suggesting that anti-proliferative therapies may be ineffective in the early post-transplant setting. However, persistent recipient HSPCs highly expressed putative targets warranting further validation. These included *PRAME*, encoding a tumor-associated antigen which may be amenable to immunotherapeutic targeting^39,40^; *CALCRL,* encoding the Calcitonin Receptor Like Receptor, a reported leukemia stem cell surface marker that has been associated with AML relapse-initiating cells^41,42^; *RXFP1*, encoding the Relaxin Family Peptide Receptor 1 that was linked to proliferation and chemoresistance in solid tumors^43,44^; and *CDKN1C*, encoding the Cyclin Dependent Kinase Inhibitor 1C or p57^Kip2^, a positive regulator of hematopoietic and leukemia stem cell quiescence that is associated with poor survival in AML/MDS after chemotherapy^45–47^. Thus, combining single-cell transcriptional and genetic profiling elucidated rare cells that may drive disease progression, and its application to patient samples helped identify potential therapeutic targets.

Overall, our results refine the understanding of immune reconstitution dynamics to identify relapse-associated features, support the distinct immune environment in *TP53*-mutated AML/MDS, and nominate immunotherapeutic targets in persistent recipient cells. These findings demonstrate the potential of dissecting differentiation states and genetic lesions directly in primary human cells to inform strategies that prevent relapse and improve post-transplant outcomes.

## RESOURCE AVAILABILITY

The following resources are provided for peer review purposes only and are confidential until publication. A complete Seurat object comprising 33 patients, 49 samples, gene expression for 496,691 cells, and annotations including donor/recipient and CNV calls has been deposited at Figshare (https://figshare.com/s/65f3672da2652943ab19). Complete code to reproduce all analyses has been provided as a zipped archive for reviewers. Single-cell sequencing data and gene expression matrices will be made available in Gene Expression Omnibus at the reopening of the US government. All data and code will be made publicly available upon publication.

## ACKNOWLEDGEMENTS

We thank the patients who participated in this study, Doreen Hearsay, the Pasquarello Tissue Bank, and the Connell and O’Reilly Families Cell Manipulation Core for providing samples, and Van Galen Lab members for helpful discussions. This work was supported by the David and Carlie Krolick Foundation and the Jock and Bunny Adams Foundation. Dr. Van Galen is supported by the National Cancer Institute Innovative Molecular Analysis Technologies (IMAT) Program (R33 CA278393), the Edward P. Evans Foundation, the Vera and Joseph Dresner Foundation, the MPN Research Foundation, an American Cancer Society Research Scholar Grant (RSG-24-1318769-01-CDP, https://doi.org/10.53354/ACS.RSG-24-1318769-01-CDP.pc.gr.222076), the Hevolution/American Federation for Aging Research (AFAR) New Investigator Award in Aging Biology and Geroscience Research, and the Krantz Family Center for Cancer Research. Drs. Van Galen and Lane are supported by the Ludwig Center at Harvard. Dr. Li is supported by NCI R50 Research Specialist Award (R50CA251956).

## DECLARATION OF GENERATIVE AI AND AI-ASSISTED TECHNOLOGIES IN THE MANUSCRIPT PREPARATION PROCESS

The authors used ChatGPT (OpenAI) and Claude (Anthropic) to assist with coding for efficiency and editing of author-written text for clarity. The authors reviewed and edited all AI-generated content and take full responsibility for its accuracy.

## METHODS

### Patient characteristics

Bone marrow aspirates were obtained from 33 patients with acute myeloid leukemia (AML) or myelodysplastic syndromes (MDS) who underwent T cell-replete allogeneic hematopoietic stem cell transplantation (HSCT) at the Dana-Farber Cancer Institute. All patients received T-cell replete transplants with tacrolimus-based prophylaxis along with methotrexate or sirolimus. One patient received post-transplant cyclophosphamide. ATG or other T-cell depletion strategies were not used in this cohort. None of the patients received DLI. Following transplantation, 7 patients in the long-term remission group and 3 patients in the relapse cohort developed acute GVHD. Patients were stratified based on post-transplant outcomes: remission (n=19), relapsed (n=10), or early relapse in the first 6 months (n=4). Clinical variables, including cytogenetics, *TP53* mutation status, conditioning regimen, and donor characteristics, were extracted from electronic medical records under institutional IRB-approved protocols and can be found in **Suppl. Table 1.** All patients provided informed consent in accordance with the Declaration of Helsinki.

### Sample collection

Bone marrow aspirates were collected as part of a routine protocol before and after HSCT at the Dana-Farber/Harvard Cancer Center Cell Manipulation Core Facility. Samples were processed on the same day and cryopreserved in liquid nitrogen storage until use.

### Sample processing and library preparation for scRNA-seq and scTCR-seq

Cryopreserved cells were thawed using standard procedures and stained with propidium iodide (PI) and CD3-FITC (BD cat. # 349201). Viable (PI-) mononuclear cells and, given sufficient cell numbers, CD3+ T cells were sorted into PBS with 1% fetal bovine serum (FBS) and counted manually to ensure accurate input for downstream applications. Single-cell gene expression and TCR libraries were generated using the Chromium Next GEM Single Cell 5’ HT Kit v2 or GEM-X 5′ Kit v3 with the Chromium Single Cell Human TCR Amplification Kit (10x Genomics) following the manufacturer’s protocol. scRNA-seq and scTCR-seq libraries were sequenced on the Illumina NovaSeq 6000 platform aiming for a read depth of 50,000 reads per cell for gene expression and 5,000 reads per cell for TCR sequencing.

### Preprocessing of single-cell RNA and TCR sequencing data

Raw sequencing data were processed with Cell Ranger (v8.0.1, 10x Genomics) for alignment to the GRCh38 reference genome, barcode filtering, and UMI quantification. TCR contig annotation was performed using the Cell Ranger V(D)J pipeline. Downstream processing was performed in R (v4.3.1) using Seurat (v5.2.1)^48^. Cells with fewer than 250 detected genes (nFeature_RNA), fewer than 500 total counts (nCount_RNA), or more than 20% mitochondrial reads were excluded as low quality.

### Cell annotation

Dimensionality reduction was performed using a standard Seurat pipeline, including principal component analysis (PCA), followed by Uniform Manifold Approximation and Projection (UMAP) for visualization. Clustering was conducted using the Louvain algorithm. Cell type annotation was performed using the reference-based label transfer BoneMarrowMap framework^25^. The `predict_CellTypes()` function was used to classify query cells and project them onto a hematopoietic reference map via K-nearest neighbor (KNN), assigning labels based on nearest neighbors. Cells that did not pass the mapping quality control filter (37,693/496,691) were excluded from downstream analyses. The algorithm provides broad and granular cell type annotations; to best represent our data, we used broad labels plus the following granular labels: CD14 Mono, CD16 Mono, Immature B, Mature B, CD4 Naive, CD4 Central Memory, CD4 Effector Memory, CD4 Regulatory, CD8 Naive, CD8 Central Memory, CD8 Effector Memory 1, CD8 Effector Memory 2, CD8 Tissue Resident Memory, T Proliferating, NK CD56high, NK Proliferating (**Figure 2A**).

### T-CellAnnoTator (TCAT)

To obtain more granular annotations of T cell subsets and identify potential differences in gene expression programs, used the starCAT package^28^. We first subsetted the complete Seurat object for T cell annotations from BoneMarrowMap (CD4 Naive, CD4 Central Memory, CD4 Effector Memory, CD4 Regulatory, CD8 Naive, CD8 Central Memory, CD8 Effector Memory 1, CD8 Effector Memory 2, CD8 Tissue Resident Memory, T Proliferating). We extracted gene expression counts and cell barcodes as input for the `starcat` algorithm with reference TCAT.V1. The algorithm’s output comprises one text file with TCAT-predicted T cell type labels and one text file with gene expression program usages. To compare the usages of 24 T cell activity programs between the long-term remission and relapse cohorts ∼3 months after transplant, we calculated the mean usage of each activity program within each cell type for each patient. To visualize these data, the median usage of programs was calculated per cell type across patients in each cohort and shown in a heatmap (**Suppl. Figure 2C**)^49,50^. To calculate significance, we used a Wilcoxon test to compare program usages in long-term remission vs. relapse patients for each cell type. After Benjamini–Hochberg correction for all 24 activity programs within each T cell type, no significant differences remained.

### Pseudotime analysis

To generate T cell trajectories, we first subsetted the Seurat object for relevant TCAT labels for CD4+ (CD4_Naive, CD4_CM, CD4_EM) or CD8+ (CD8_Naive, CD8_CM, CD8_EM, CD8_TEMRA) cells, and excluded proliferating T cells. We then ran dimensionality reduction on the subsetted CD4+ or CD8+ T cells objects using standard Seurat functions, removing batch effects with Harmony (**Suppl. Figure 2D-E**)^51^. Next, we used the Monocle3 workflow^52^ with default parameters except `learn_graph` with `use_partition = F` because data was already subsetted to closely related T cells. Naïve T cells were used as a root, and Monocle3 pseudotime values were added as a metadata column to the (subsetted) Seurat object. Finally, to ensure that results were not driven by differences in cell numbers, an equal number of cells was sampled from each patient (913 CD4+ T cells or 434 CD8+ T cells per patient), and ggplot’s `geom_density` function was used to create density plots with Monocle3 pseudotime values (**Figure 2E-F**). For statistics, we calculated the proportion of cells that fell within three pseudotime bins—low, medium, and high—for each patient. These proportions were compared between patients in the long-term remission and relapse cohorts (**Suppl. Figure 2F**) using a Wilcoxon test.

For malignant cells, pseudotime values from the BoneMarrowMap framework were derived using the `predict_Pseudotime()` function, which assigns each cell a position along the hematopoietic differentiation hierarchy. Cells were subsetted for Numbat’s “tumor” compartment, for myeloid/erythroid lineages (HSC MPP, MEP, EoBasoMast Precursor, Megakaryocyte Precursor, Early Erythroid, Late Erythroid, LMPP, Cycling Progenitor, Early GMP, Late GMP, Pro-Monocyte, CD14 Mono, CD16 Mono, cDC, pDC), and for patients with both pre-transplant and relapse malignant cells (P20, P23, P30, P31, **Suppl. Figure 6A**). Then, data was subsetted to the same number of cells per patient/status (n=127 cells) to account for different cell numbers, and cell densities were plotted using `geom_density()`. *P*-values were calculated using the Kolmogorov–Smirnov test either on all patients together (**Figure 5D**) or the individual patients (**Suppl. Figure 6F**).

### TCR analysis

T cell receptor (TCR) α and β chain sequences were extracted using the scRepertoire package (v2.0)^53^. Clonotypes were defined using the `CTStrict` parameter, which considers identical V(D)J gene usage and CDR3 nucleotide sequences across paired TCR chains. We then sampled 300 T cells per sample, and excluded samples with fewer than 300 T cells. TCR diversity was quantified using the inverse Simpson index, defined as:

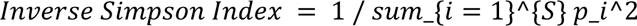

where S is the total number of clonotypes and p is the proportional abundance of clonotype within a given patient.

### Differential gene expression and gene set enrichment analysis

For differential gene expression analysis, we used Seurat’s `AggregateExpression` function to pseudobulk cells of interest by patient (relapse vs. pre-transplant malignant cells in **Figure 5** and **Suppl. Figure 6**, donor vs. recipient HSPCs in **Figure 6** and **Suppl. Figure 7**). We then used DESeq2 to identify differentially expressed genes and shrunk log fold changes using the apeglm method^54,55^. Genes were ranked by log2 fold change for Fast Gene Set Enrichment Analysis (FGSEA)^56^ using Hallmark (**Figure 6C**) or curated C2 (**Suppl. Figure 6C**) pathways^57^.

### Souporcell analysis

Donor and recipient chimerism was resolved using Souporcell (v2.4), a genotype-free Bayesian clustering algorithm for single-cell RNA-seq data^22^. BAM files generated by Cell Ranger were used as input. First, different time points from the same patient were merged using `samtools merge`. The resulting merged BAM files were processed using the default panel of common single-nucleotide polymorphisms (SNPs) provided by Souporcell. The analysis was conducted using Souporcell’s Docker container, with the number of genotype clusters specified as `--k 2` to represent recipient and donor populations. Cells were assigned to genotype clusters based on posterior probability, and only those with assignment probability ≥0.9 were retained. Doublets and ambiguous cells were excluded from downstream analyses.

To assign genotype clusters to donor or recipient identity in the absence of matched germline reference genotypes, we developed a rule-based classification script. When available, pre-transplant samples were used to identify the recipient genotype as the one comprising >80% of cells. In samples lacking a pre-transplant reference, the genotype with >80% representation across all cell types was classified as donor. Patients lacking a clear majority genotype (12/33 patients) were labeled as unknown and excluded from downstream analyses. Using these rules, we successfully resolved cell-of-origin in 21 of 33 patients, yielding donor or recipient assignments for 342,780 out of 496,691 total cells.

### Copy number analysis with Numbat

To infer somatic copy number variation (CNVs) and clonal architecture, we used Numbat (v0.5.2), a haplotype-aware probabilistic framework that integrates gene expression, allelic imbalance, and population-derived haplotypes to detect allele-specific CNVs and infer lineage relationships from single-cell RNA-seq data^23^. Following donor–recipient deconvolution with Souporcell, we identified 155,392 recipient-derived cells across multiple time points from 21 patients. For each patient, recipient cell barcodes were used to subset the corresponding BAM files with the 10x Genomics subset-bam package. Preprocessing was carried out on a virtual machine instance using the Numbat Docker container, where the `pileup_and_phase.R` script was used to perform SNP pileup, phasing, and allele counting with the 1000 Genomes Phase 3 SNP VCF (hg38), a genetic map, and a phased reference panel. Samples from different time points belonging to the same patient were processed together in a single run, resulting in one allele count file per time point. These allele count matrices were merged, UMI count matrices for each patient were extracted from the corresponding Seurat object, and final CNV inference was performed using Numbat’s R function `run_numbat`. CNVs were visualized as genome-wide heatmaps, and from 6 patients were classified as normal (n=21,317) or tumor (n=36,837) based on the presence of patient-specific alterations. For the remaining 15 patients, no CNVs remained after filtering based on log-likelihood ratio (LLR) thresholds in pseudobulk data (n=10) or based on entropy thresholds in single-cell data (n=2), or results did not match clinical cytogenetics (n=3). Numbat-inferred CNAs showed concordance with available clinical karyotype data (**Figure 5A, Supplemental Figure 5**), supporting the validity of this approach in characterizing clonal architecture in AML.

## SUPPLEMENTAL FIGURES

**Supplemental Figure 1.**
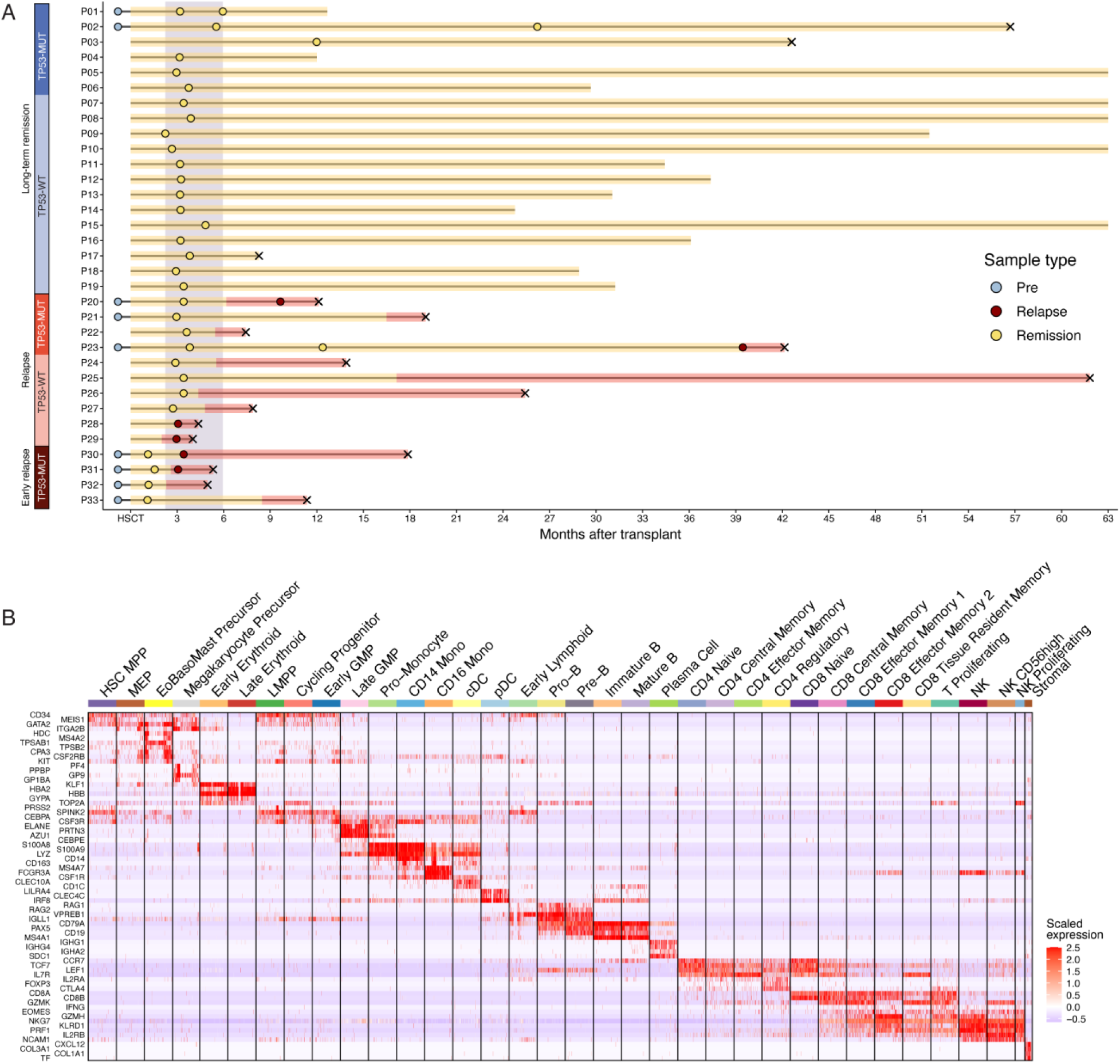
Cohort design and cell type annotation. **A.** Swimmer plot shows the clinical course of patients we analyzed. Each horizontal bar represents one patient’s timeline, ordered by cohort and *TP53* status. Circles mark time points of sample collection. Bars shaded yellow indicate clinically remission period, red indicate relapse intervals. Black × denotes death. The purple shaded region marks the remission samples on which most analyses are focused. Patients in the third cohort (early relapse) had *TP53*-mutated AML and measurable blasts at transplant; they were only used for relapse analysis in **Figure 5** and **Suppl. Figure 6**. **B.** Heatmap shows expression of cell type-specific genes (rows) across cells (columns) grouped by cell types. Many of these genes, identified using Seurat’s FindMarkers function, are well-established cell type marker genes, supporting the BoneMarrowMap annotations.

**Supplemental Figure 2.**
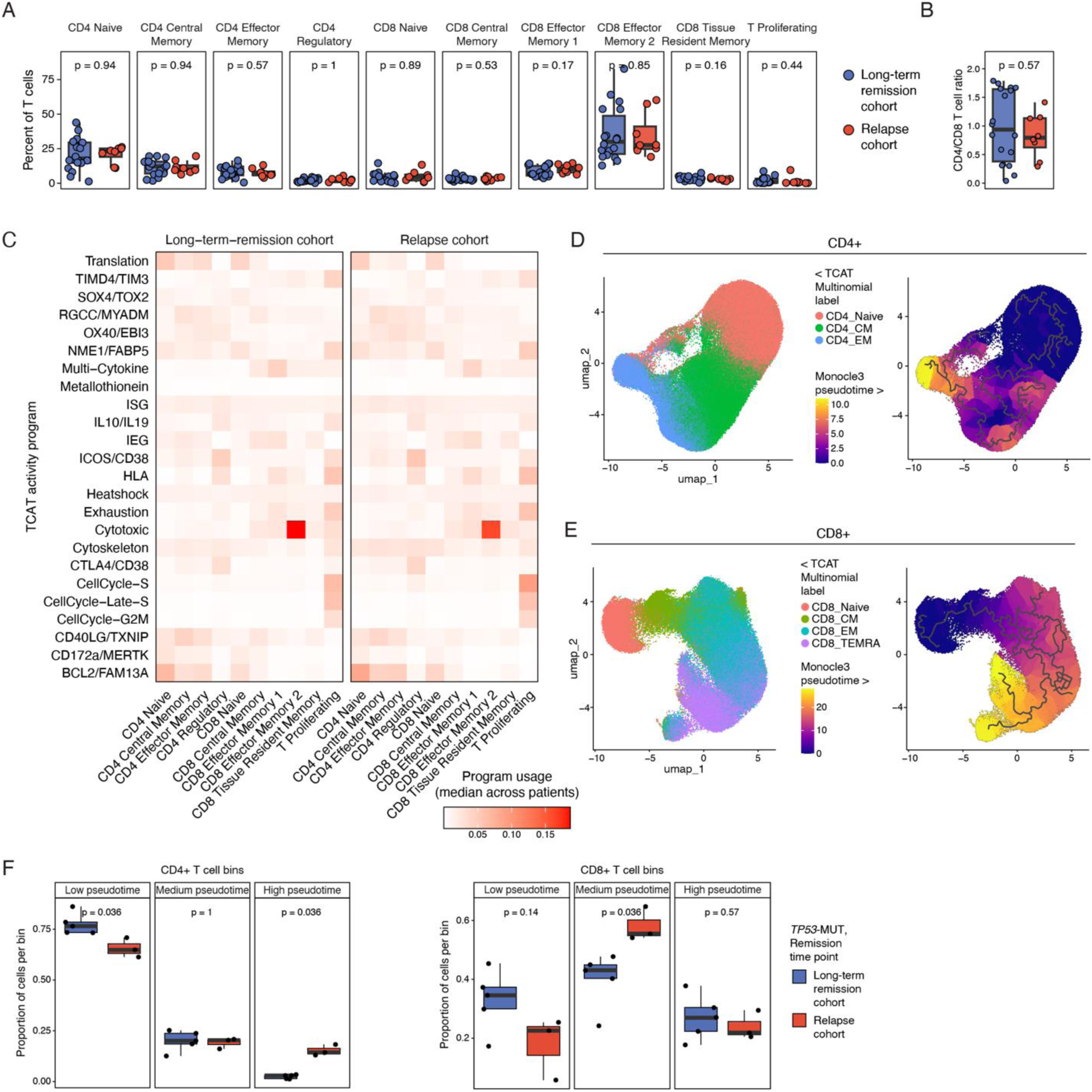
Supporting data for T cell analyses. **A.** Boxplots comparing the proportion of T cell subsets between long-term remission and relapse cohorts among all patients ∼3 months after transplant. No significant differences were detected (Wilcoxon test). **B.** Comparison of CD4/CD8 ratio between long-term remission and relapse cohorts among all patients from ∼3 months after transplant. P-value was calculated using Wilcoxon test. **C.** Heatmap shows usage of T cell activity programs (rows, defined by TCAT^28^) in T cell types (columns), separated by cohort. Each square shows median usage across patients (n=18 in the long-term-remission cohort and n=8 in the relapse cohort). Wilcoxon tests with Benjamini–Hochberg correction showed that none of the program usages differed significantly between cohorts (see **Methods**). **D-E.** UMAPs show (D) CD4+ and (E) CD8+ T cells colored by TCAT label (left) and Monocle3 pseudotime (right). The pseudotime provides a proxy for progressive differentiation states used in **Figure 2E**. Trajectory analysis was performed after subsetting CD4+ T cells to non-proliferating cells with TCAT labels CD4_Naive, CD4_CM, and CD4_EM (excluding Treg), CD8+ T cells were subsetted to non-proliferating cells with TCAT labels CD8_Naive, CD8_CM, CD8_EM, and CD8_TEMRA (excluding MAIT and gdT), and UMAP coordinates were calculated using standard Seurat functions. **F.** Boxplot shows the proportion of T cells in three pseudotime bins for patients (symbols) in the long-term remission and relapse cohorts. These analyses (like **Figure 2C-E**) are subsetted for remission samples ∼3 months after transplant from patients with *TP53*-mutated AML. For each patient, the sum of proportions adds up to 1. P-values were calculated using the Wilcoxon test.

**Supplemental Figure 3.**
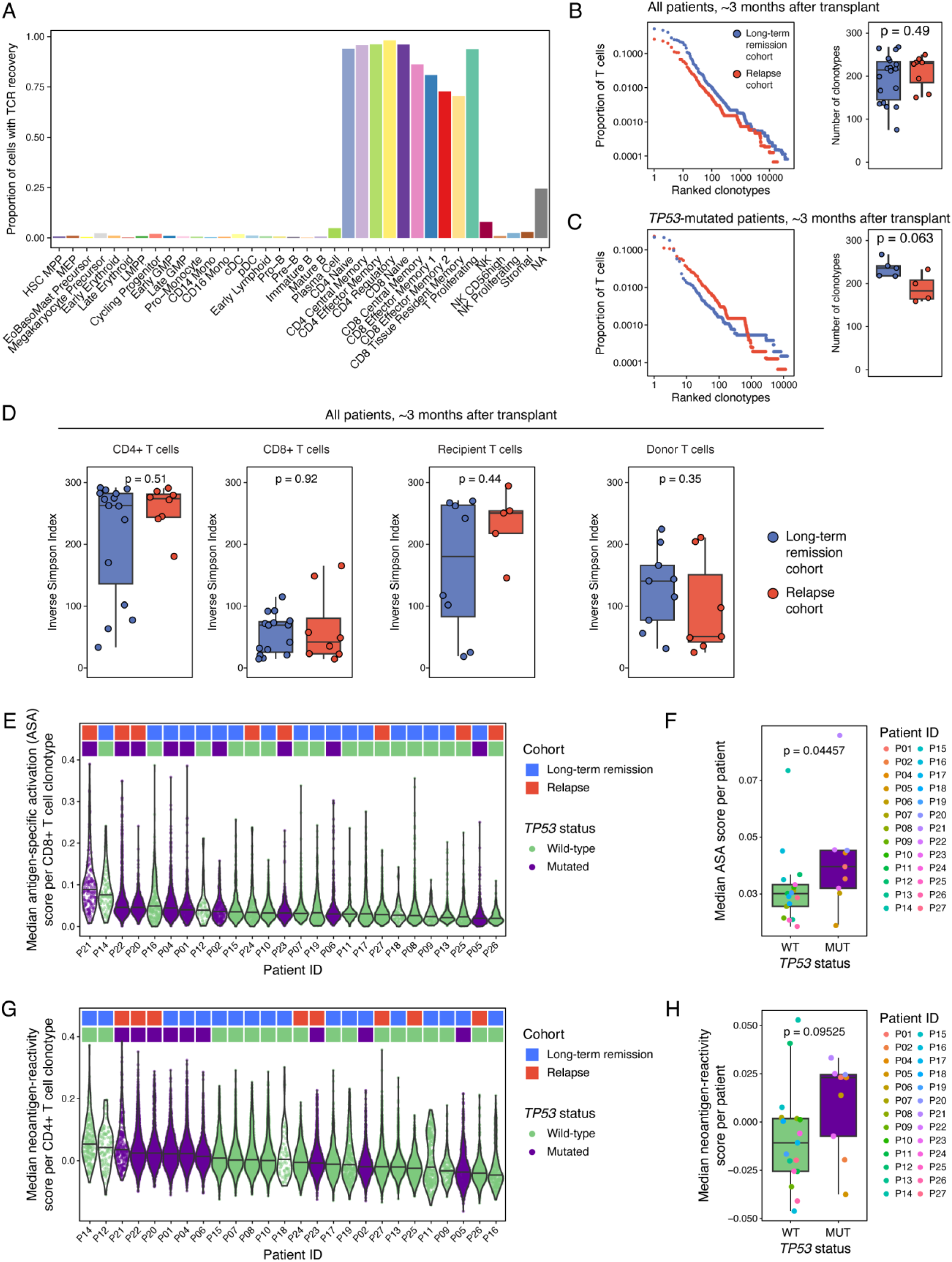
TCR recovery and analysis. **A.** Bar plot shows the proportion of cells in which a TCR was recovered across all annotated cell types. **B.** Left: Scatter plot shows the proportion of the T cell compartment that clonotypes (symbols) comprise. Clonotypes are ranked from large to small. Right: Boxplot shows the number of T cell clones per patient, separated into long-term remission (blue) and relapse (red) cohorts. Each symbol represents a patient, and data was subsetted to 300 T cells per patient. P-value was calculated using a Wilcoxon test. **C.** Same as (B), but subsetted for patients with *TP53-*mutated AML. **D.** Boxplots show TCR diversity indices ∼3 months after transplant from all patients, separated into long-term remission and relapse cohorts. Data is stratified by T cell subset (CD4+ or CD8+), and by recipient vs. donor origin. Each sample was subsetted to 300 T cells or excluded if fewer T cells were detected. P-values were calculated using Wilcoxon tests. **E.** Violin plot shows CD8+ neoantigen-specific activation scores (ASA) in CD8+ T cells ∼3 months after transplant. Each symbol represents a T cell clonotype, with values indicating the median ASA score across all cells within that clonotype. **F.** Boxplot shows median ASA scores^28^ in CD8+ T cells ∼3 months after transplant per patient, separated by *TP53* status. Each symbol represents the median score of all T cell clonotypes within a patient. P-value was calculated using Wilcoxon test. **G.** Violin plot shows CD4+ neoantigen-reactivity scores in CD4+ T cells ∼3 months after transplant. Each symbol represents a T cell clonotype, with values indicating the median NeoTCR4 score^29^ across all cells within that clonotype. **H.** Boxplot shows median CD4+ neoantigen-reactivity scores in CD4+ T cells ∼3 months after transplant per patient, separated by *TP53* status. Each symbol represents the median score of all T cell clonotypes within one patient. P-value was calculated using Wilcoxon test.

**Supplemental Figure 4.**
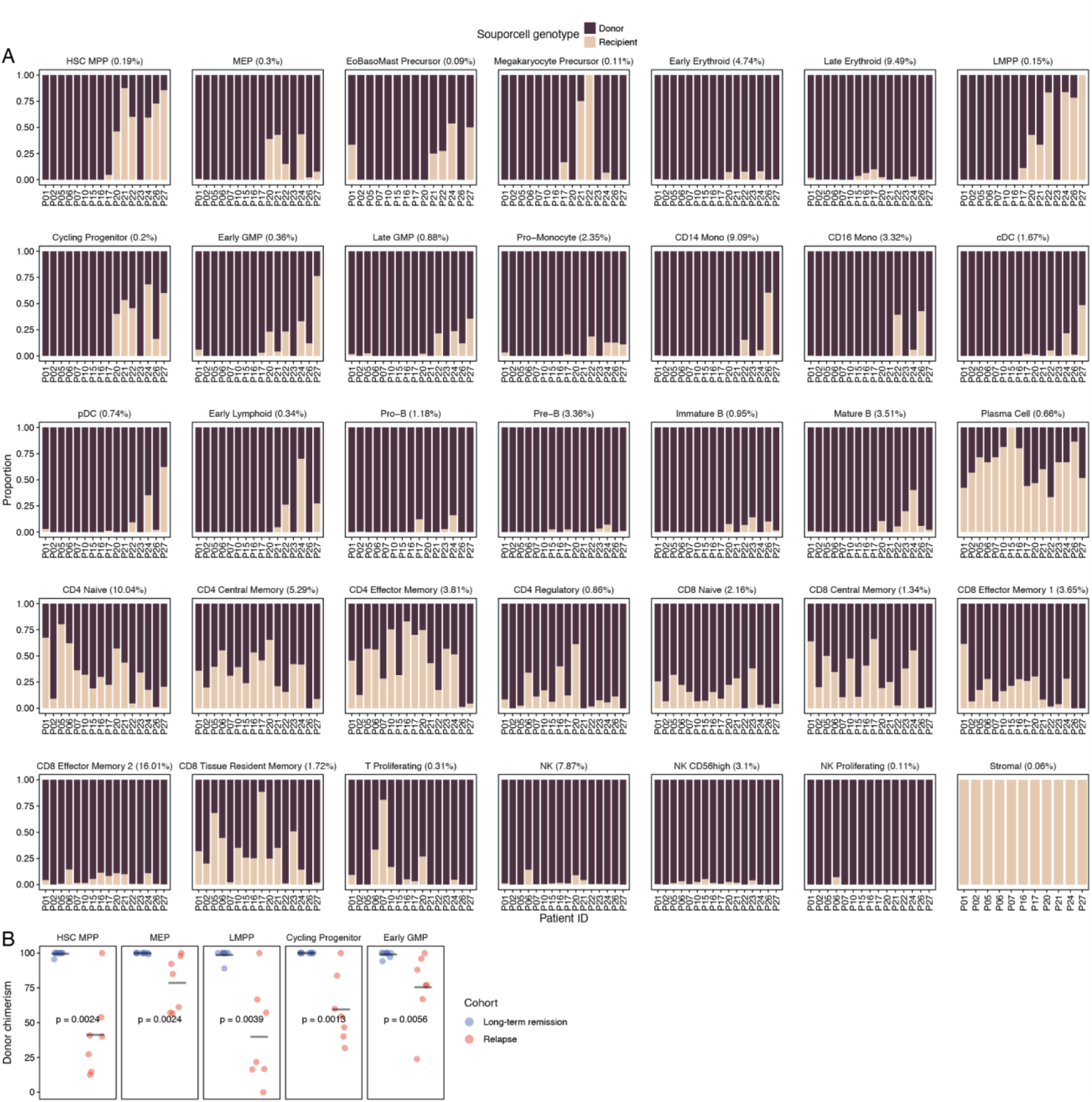
Donor and recipient cell proportions. **A.** Barplots show the proportions of recipient and donor cells (y) for each patient (x) and cell type (plots). Recipient and donor cells were classified using Souporcell, and only remission samples ∼3 months after transplant are used to generate these barplots. **B.** Donor chimerism determined by Souporcell for individual cell populations comprising **Figure 4D**: HSC/MPPs, MEPs, LMPPs, cycling progenitor cells, and early GMPs, ∼3 months after transplant.

**Supplemental Figure 5.**
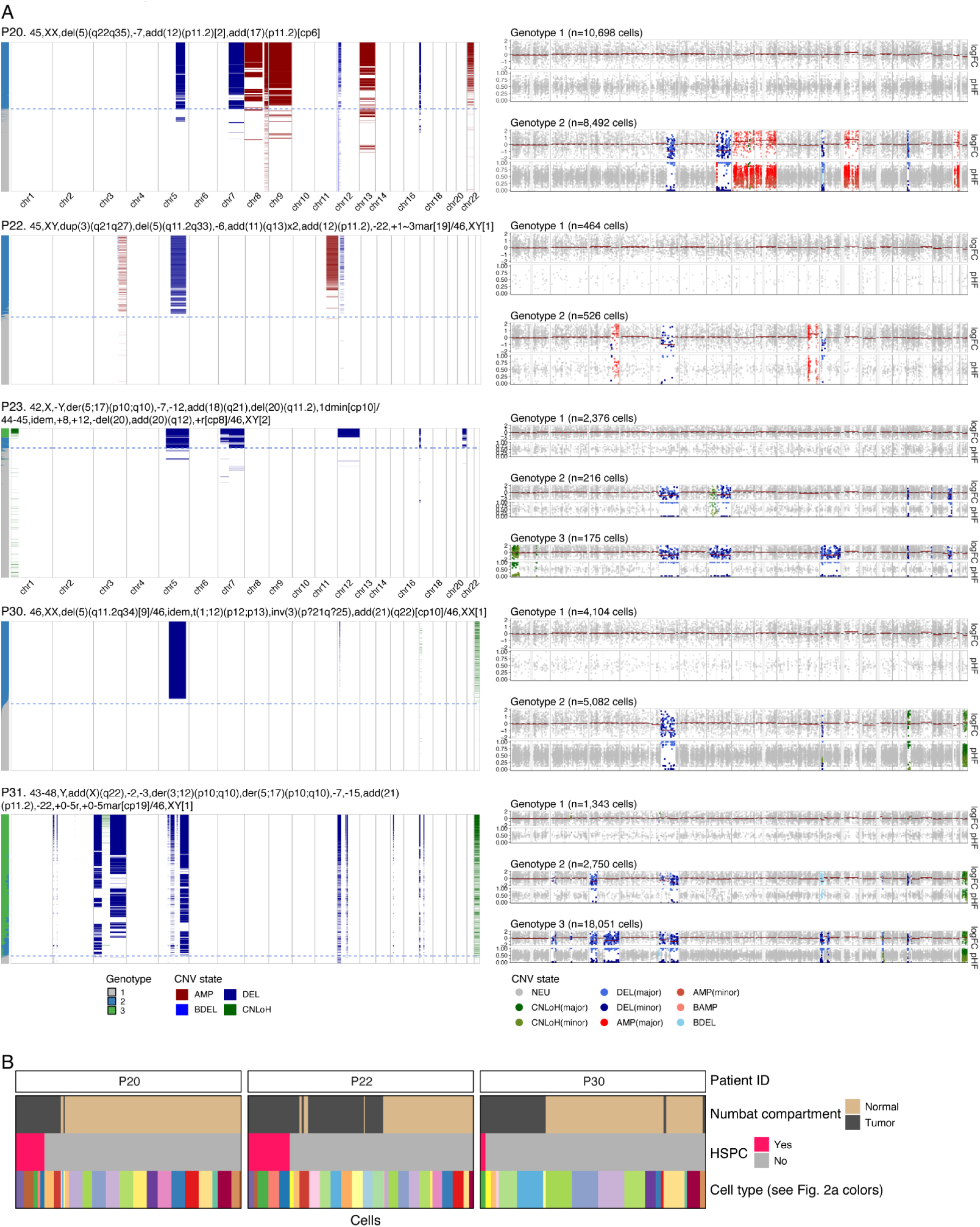
Detection of CNVs based on single-cell RNA-seq using Numbat. **A.** Left: heatmaps show genome-wide CNVs (colors) in single cells (rows), generated using Numbat. Each heatmap represents one patient, and cytogenetics determined by karyotyping are indicated above the plot. Each row represents a cell, and columns correspond to genomic positions across chromosomes 1–22. CNV states are color-coded as indicated below the heatmaps: amplifications (AMP, dark red), deletions (DEL, dark blue), biallelic deletions (BDEL, light blue), and copy-neutral loss of heterozygosity (CNLoH, green). On the left, cells are grouped by genotype: normal cells (grey) and malignant cells (blue or green). The horizontal dashed line separates malignant from normal cell clusters based on inferred CNV profiles. Right: genome plots show CNV profiles for pseudobulked clones, with pseudobulked normal cells on top (genotype 1) and pseudobulked malignant cells on the bottom. Each symbol represents a genomic bin, with read coverage (logFC, −2 to +2) shown on top and posterior haplotype frequencies (pHF, 0 to 1) on the bottom. **B.** Tile plot shows Numbat compartment (top), whether each cell is an HSPC (middle), and cell type annotation (bottom) for cells from patients in remission (columns). “HSPC” comprises the BoneMarrowMap annotations HSC MPP, MEP, LMPP, Cycling Progenitor, and Early GMP. A maximum of 10 cells per cell type is shown to improve visibility. All remission samples are included, and samples with 0 recipient HSPCs are excluded (P23, P31, P33).

**Supplemental Figure 6.**
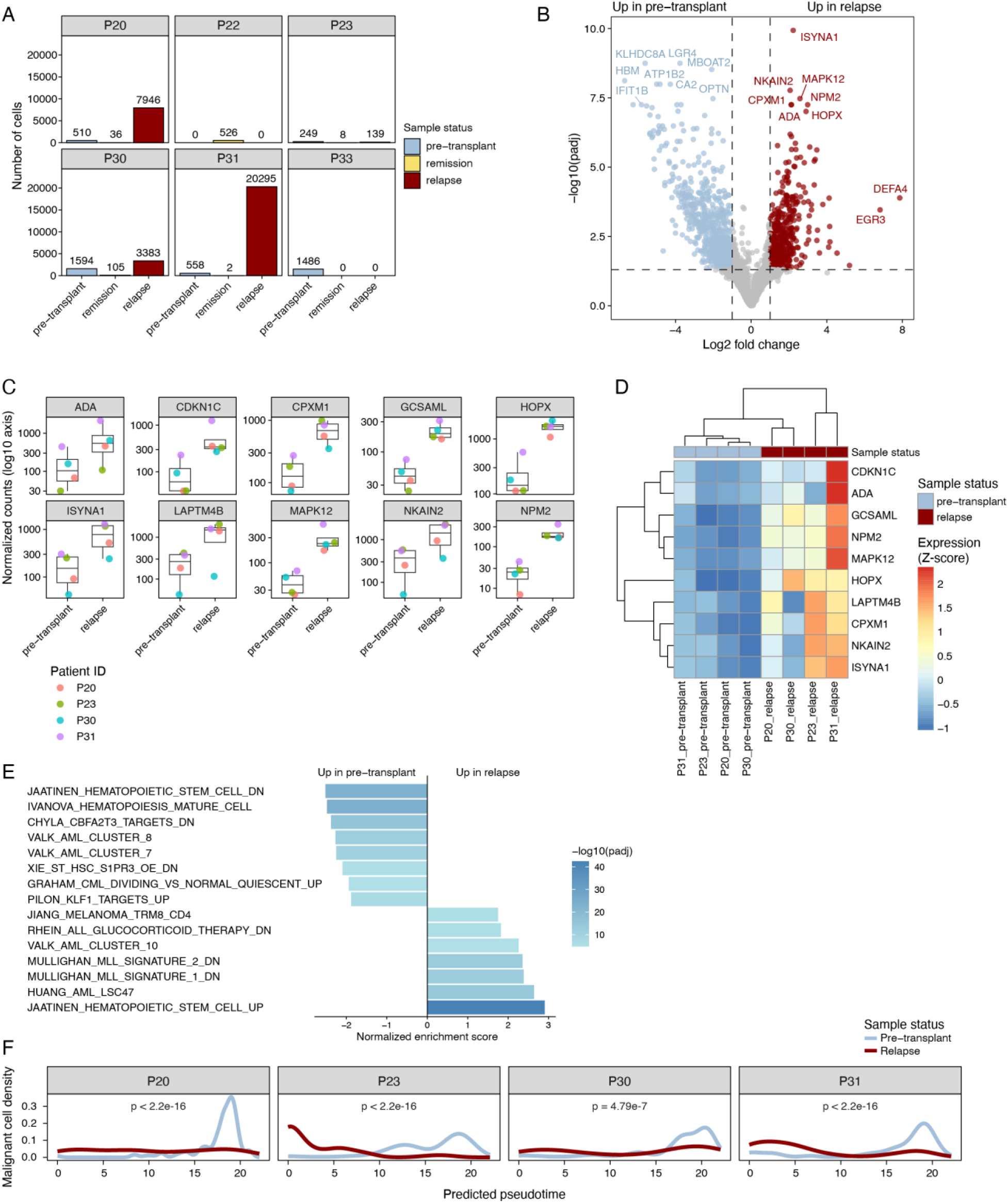
Differential gene expression of malignant cells pre-transplant vs. relapse. **A.** Barplots show the number of malignant cells in six patients where Numbat successfully ran, separated by disease status. P22 and P33 were excluded from further analysis of tumor evolution since malignant cells were only detected at one time point. Downstream differential gene expression analysis was performed after pseudobulking to account for different cell numbers. **B.** Volcano plot shows differentially expressed genes (symbols) between pre-transplant malignant cells (n=31,763 cells across four patients) and relapse malignant cells (n=2,911 cells across the same four patients). **C.** Boxplots show DESeq2-normalized counts of the top 10 genes (by adjusted p-value) upregulated in malignant cells at relapse compared to pre-transplant. Patient ID is indicated by symbol color. **D.** Heatmap shows expression of the same top 10 genes (rows) by sample (columns). Normalized expression values were z-score scaled by rows (per gene across samples). **E.** Bar plot shows top 15 pathways (by adjusted p-value) enriched in pre-transplant (left) or relapse (right) samples by GSEA. **F.** Density plot shows the distribution of predicted pseudotime values for malignant cells from pre-transplant and relapse samples, separated by patient. Pseudotime values were determined by mapping cells onto the BoneMarrowMap reference. In relapse samples, the proportion of primitive leukemia cells (left) is higher while the proportion of differentiated cells (right) is lower. Statistical significance was assessed using the Kolmogorov–Smirnov test.

**Supplemental Figure 7.**
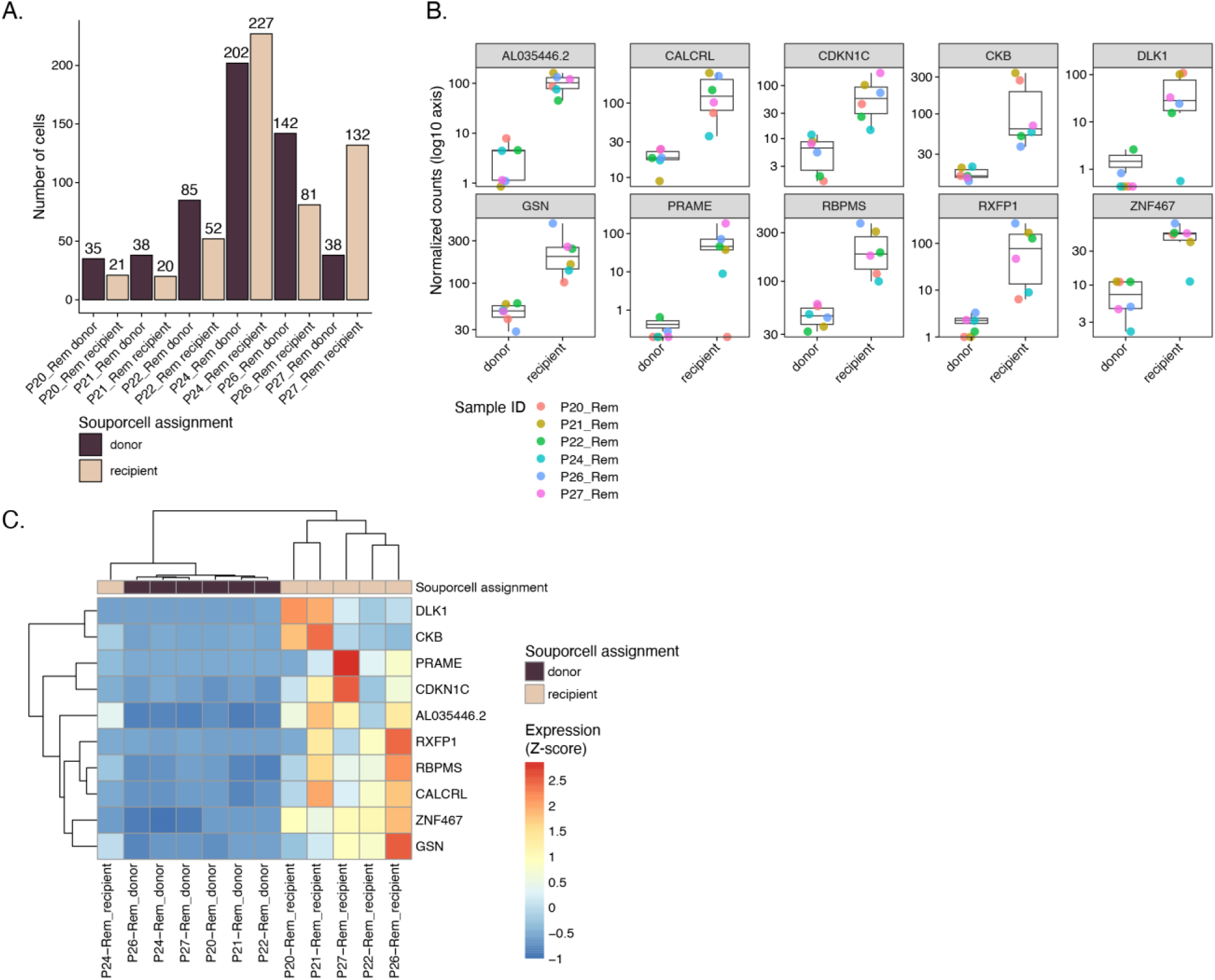
Differential gene expression of recipient vs. donor HSPCs. **A.** Bar plot shows the number of HSPCs (y) in six patients, separated by Souporcell assignment. All six patients are from the relapse cohort and cells are subsetted to remission samples ∼3 months after transplant. X-axis label indicates patient, disease status (remission), and Souporcell assignment. Downstream differential gene expression analysis was performed after pseudobulking to account for different cell numbers. **B.** Boxplots show DESeq2-normalized counts of the top 10 genes (by adjusted p-value) upregulated in recipient HSPCs compared to donor HSPCs in remission. Patient ID is indicated by symbol color. **C.** Heatmap shows expression of the same top 10 genes (rows) by sample (columns). Normalized expression values were z-score scaled by rows (per gene across samples).

## SUPPLEMENTAL TABLES

**Supplemental Table 1:** All patient information.

**Supplemental Table 2:** Differentially expressed genes and pathways between malignant cells at the pre-transplant and relapse time points (corresponding to **Suppl. Figure 6B and E**). Genes with a positive log fold change and pathways with a positive enrichment score are increased at relapse.

**Supplemental Table 3:** Differentially expressed genes and pathways between recipient HSPCs and donor HSPCs ∼3 months after transplant in remission samples (corresponding to **Figure 6B-C**). Genes with a positive log fold change and pathways with a positive enrichment score are increased in recipient HSPCs.

